# Structural Basis for SARS-CoV-2 Envelope Protein in Recognition of Human Cell Junction Protein PALS1

**DOI:** 10.1101/2021.02.22.432373

**Authors:** Jin Chai, Yuanheng Cai, Changxu Pang, Liguo Wang, Sean McSweeney, John Shanklin, Qun Liu

## Abstract

The COVID-19 pandemic caused by the SARS-CoV-2 virus has created a global health and economic emergency. SARS-CoV-2 viruses hijack human proteins to promote their spread and virulence including the interactions involving the viral envelope (E) protein and human proteins. To understand the structural basis for SARS-CoV-2 viral-host recognition, we used cryo-electron microscopy to determine a structure for the human cell junction protein PALS1 and SARS-CoV-2 E protein complex. The structure shows that the E protein C-terminal DLLV motif recognizes a pocket formed exclusively by hydrophobic residues from the PDZ and SH3 domains in PALS1. Our structural analysis provides an explanation for the observation that the viral E protein recruits PALS1 from lung epithelial cell junctions resulting in vascular leakage, lung damage, viral spread, and virulence. In addition, our structure provides novel targets for peptide- and small-molecule inhibitors that could block the PALS1-E interactions to reduce the E-mediated damage to vascular structures.

## Main

Severe acute respiratory syndrome coronavirus 2 (SARS-CoV-2) is a causative agent for the COVID-19 pandemic that is disrupting human health and global economy. Although 80% of COVID-19 patients display mild or no symptoms, 20% of them developed serious conditions mostly in the population of elderly person and those with underlying pre-existing medical conditions. The virus has caused more than two million deaths and more than 100 million cases worldwide. Most deaths are associated with an acute respiratory distress syndrome (ARDS) and tissue damage linked to virus-induced hyper-immune responses ^1^.

SARS-CoV-2 and SARS-CoV-1 genomes encode a small envelope (E) protein that is a critically important component in viral life cycle of assembly, release, and virulence ^2^. SARS-CoV-2 E is composed of 75 amino acid residues with two distinct domains: an N-terminal transmembrane (TM) domain followed by a C-terminal domain. E is a multifunctional protein. Besides its structural roles required to induce membrane curvature for viral assembly in cooperation with the viral membrane (M) protein, E mediates host immune responses through two distinct mechanisms: a pore-forming TM domain related to the activation of NLRP3-inflammasome ^3^; and a PDZ (PSD-95/Dlg/ZO-1)-binding function via its C-terminal domain ^4,5^. Structurally, the TM domain of SARS-CoV-2 E forms a pentameric ion channel, similar to that of SARS-CoV-1 E ^6,7^. However, the C-terminal domain has no well-defined structure, perhaps due to the lack of a stable complex.

In humans, there are about 150 unique proteins encoding one or more PDZ domains. These PDZ domains contain 80-110 amino acid residues and are essential in regulating human immune responses and numerous physiological and pathological activities ^8^. PDZ-domain-containing proteins in cell junctions have been hijacked by various viruses to potentiate their virulence ^8^. Both SARS-CoV-2 and SARS-CoV-1 E proteins harbor a PDZ-binding motif (PBM) at their C-termini. Although the exact mechanism is unknown, interactions between the PBM and a human cell junction protein, PALS1, showed that E causes the relocation of PALS1 from the cell junction to the endoplasmic-reticulum–Golgi intermediate compartment (ERGIC) site where E is localized and virus assembly and maturation occur ^4^. As a cell junction protein, PALS1 is a structural component of an apical Crumbs (Crb) complex in the establishment and maintenance of cell polarity and intercellular tight junctions ^9,10^. In addition to PALS1, E also interacts to PDZ-containing adhesion junction protein syntenin ^5^, tight junction protein ZO1 ^11^ and other cell-junction proteins ^12^. The relocation of these cell-junction proteins in lung epithelial cells contributes to vascular leakage, diffuse alveolar damage (DAD), cytokine storm initiation, and acute respiratory distress syndrome (ARDS), commonly leading to death in elderly COVID-19 patients and those with underlying conditions ^2^.

The PBM in E contains four conserved residues (DLLV) and is conserved between SARS-CoV-2 and SARS-CoV-1 viruses. The motif appears to play a critical role in virulence because mutants without the PBM are either attenuated or nonviable ^5,13^. Binding assays using the 10-residue C-terminal peptides of SARS-CoV-2 E and SARS-CoV-1 E show enhanced binding affinity of SARS-CoV-2 E peptide to the PDZ domain in PALS1 ^14^. However, there is a lack of structural information to define such protein-protein interactions, which hinders further understanding of the mechanisms of the E-mediated virulence. In this work, we describe the structure of the PALS1-E complex to define the mechanism of recognition of the PALS1 PDZ and SH3 domains by the C-terminal PBM of the E protein.

## Results

### Production of the PALS1-E complex

PALS1 contains five domains, two N-terminal L27 domains and three C-terminal domains, PDZ, SH3, and GK (named as PSG). To improve protein stability, we expressed and purified the PSG (residues 236–675) without a loop between the SH3 and GK domains ^9^ (**Fig. S1a**). The expressed protein was purified by Ni-NTA affinity resins followed by size-exclusion chromatography (SEC) (**Fig. S1b**). Based on the SEC analysis, we found that majority of PSG is a dimer (**Fig. S1c**).

To study the structural basis for recognition of PALS1 PSG by SARS-CoV-2 E C-term domain, we synthesized an E C-term 18-amino-acid peptide (Ec18) containing the PBM (**Fig. 1a**). To check the biding affinity between Ec18 and PSG, we labeled PSG using a fluorescence dye and titrated it using a serial dilution of Ec18. Using a microscale thermophoresis method ^15^, we measured the K*d* as 11.2 μM (**Fig. S1d**). Our measured value is consistent with the binding affinity using a 10-aa peptide and the PDZ domain alone; where the measured K*d* is 40 μm (Toto et al., 2020). Considering the low affinity between Ec18 and PSG, we used a high ratio of Ec18 for complex formation by incubating purified PSG with Ec18 at a molar ratio of 1:10 for 2hrs at room temperature.

**Figure 1.**
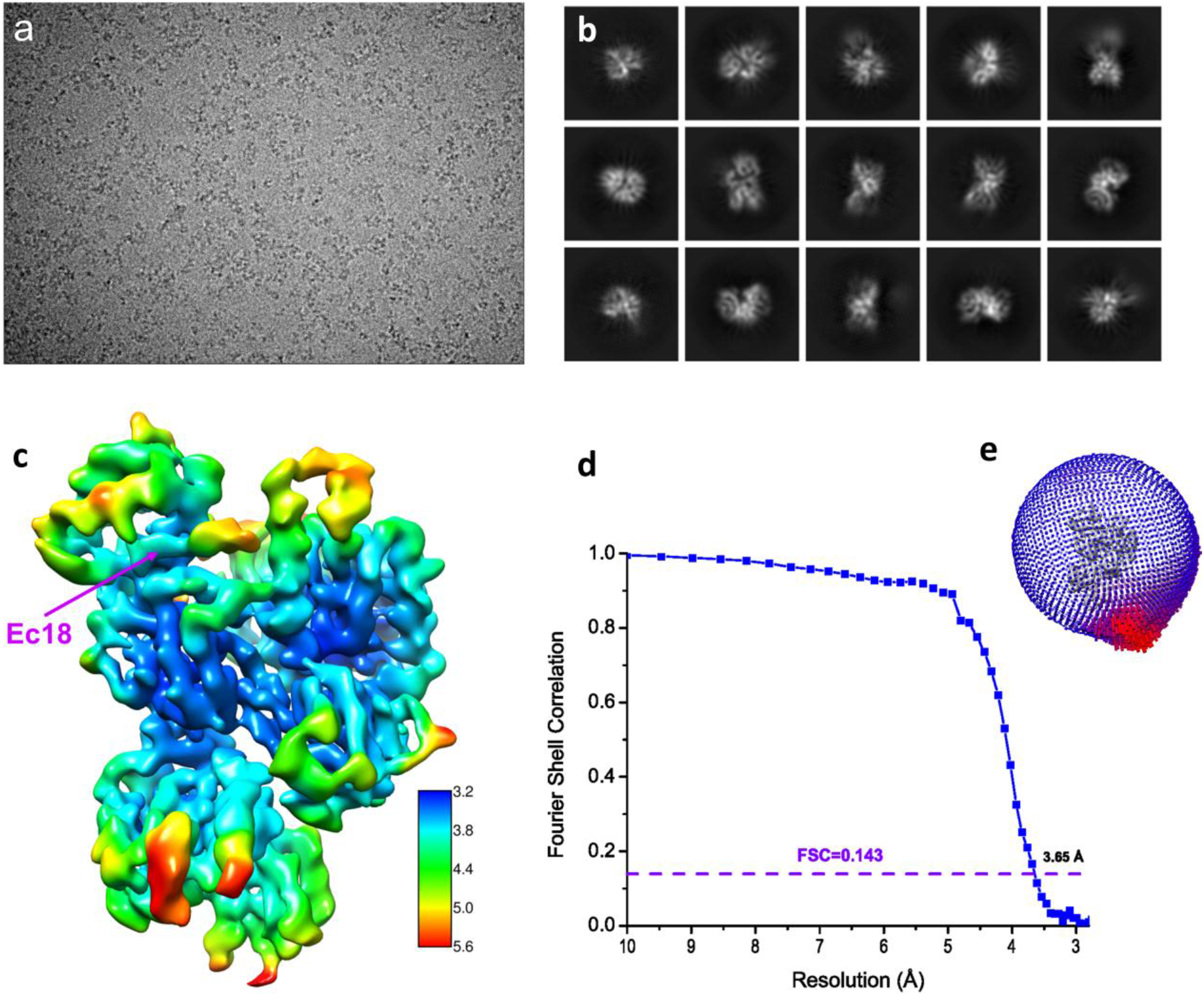
Structure determination by single-particle Cryo-EM. (**a**) A typical motion-corrected Cryo-EM micrograph. (**b**) 2D class averages. (**c**) Reconstructed map colored with local resolutions. (**d**) Fourier Shell Correlation (FSC) curve for the 3D reconstruction to determine the structure resolution. (**e**) Orientation distribution for particles used in 3D reconstruction.

### Structure determination

We subjected the PSG-Ec18 complex to analysis using single-particle Cryo-EM. Our initial 2D class averages showed a preferred particle orientation. To get additional views of the complex, we performed detergent screening and found that the inclusion of 0.05% CHAPS allowed PSG-Ec18 particles to distribute evenly (**Fig. 1a**) and helped us obtain multiple views after 2D class averaging (**Fig. 1b, 1e**). We optimized our particle picking procedure using a local dynamic mask for defocus-based particle picking ^16^. After iterative 2D and 3D classifications and refinements with per-particle CTF and Bayesian polishing (**Fig. S2**) with Relion3 and CryoSPARC ^17,18^, we obtained a final reconstruction at 3.65 Å (**Fig. 1c**) using Fourier Shell Correlation of 0.143 as a cutoff (**Fig. 1d**). The map shows clear secondary structures and side chains that allowed us to build and refine atomic models (**Fig. S3**).

### Structure of the PSG-Ec18 complex

The solved structure contains a dimer of PSG and a single Ec18 (**Fig. 2a**). In one PSG monomer, the PDZ, SH3, and GK domains were observed; while in the other monomer, the PDZ domain was missing. However, in our initial 3D classification, we observed a class with a highly disordered region corresponding to the missing PDZ domain (**Fig. S2**). In comparison to the crystal structure of the PSG dimer in complex with its physiological ligand Crb-CT ^9^, the PDZ and SH3 domains are rotated about 38° relative to the GK domain in the PSG-Ec18 complex (**Fig. S4a**).

**Figure 2.**
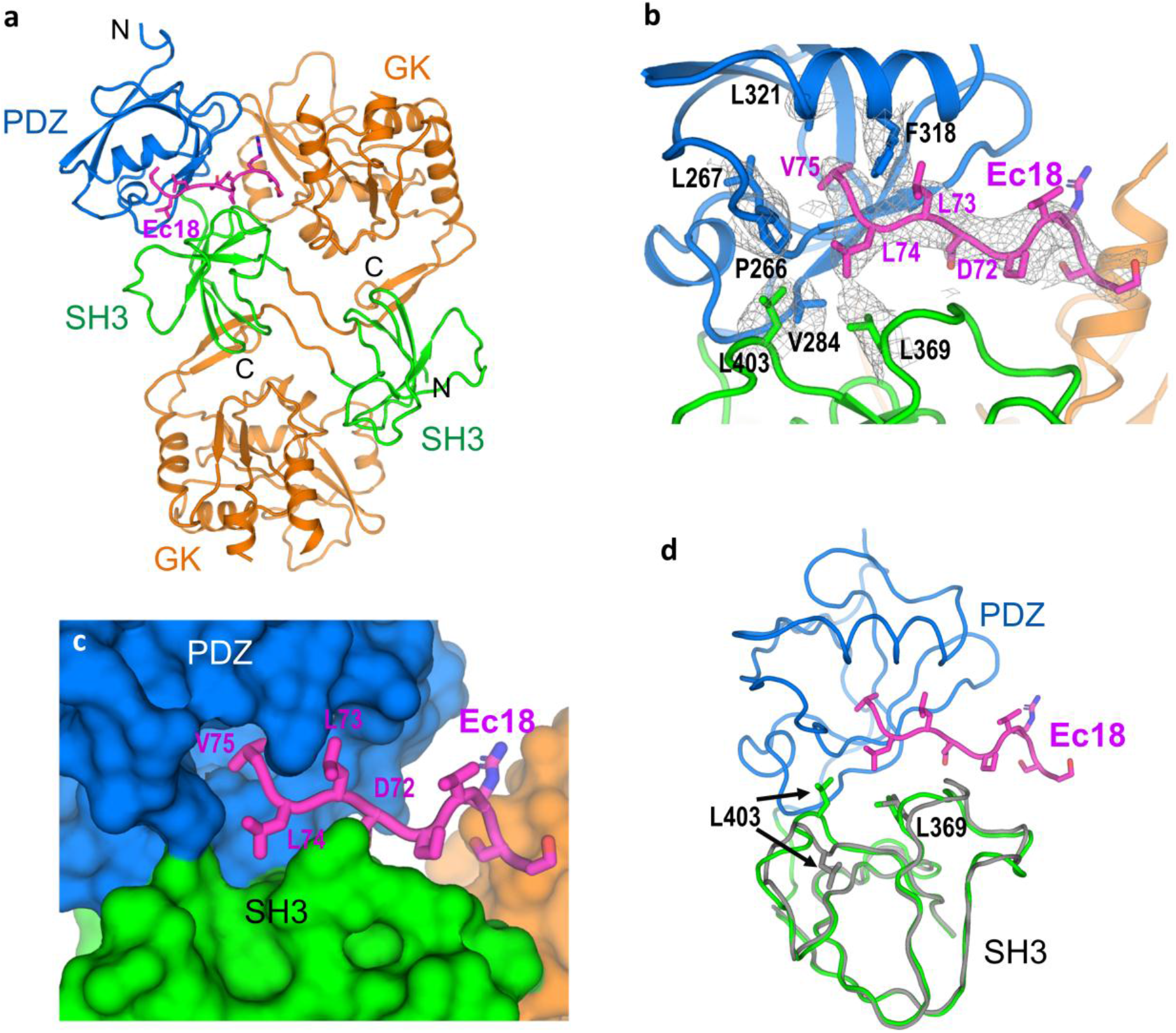
Recognition of the PALS1 PDZ and SH3 domains by the E PBM. (**a**) Structure of the PSG-Ec18 complex shown as cartoons with different colors for different domains. Ec18 is shown as magenta sticks. (**b**) Binding site structure. Hydrophobic residues forming the binding pocket were shown as sticks. Potential density map for the binding site is shown as gray isomeshes contoured at 5.5σ. (**c**) Surface representation of the binding site with Ec18 showing as sticks. (**d**) Superimposition of the two PSG monomers to show conformational changes in SH3 domains. The PDZ domain in the second monomer is disordered.

In our structure, Ec18 is inserted in a hydrophobic pocket between the PDZ and SH3 domains through the PBM(72DLLV75) (**Fig. 2b, c**). The density coverage for residues Leu74-Val75 on the Ec18 and Phe318 on the PDZ domain are well defined and help position Ec18 in the binding pocket. Residues Phe318, Leu321, Leu267, Pro266, and Val284 from PDZ and Leu369 and Leu403 from SH3 are involved in forming the hydrophobic binding pocket. Phe318, Leu369, and Leu403 in PALS1 and Leu74 and Val75 in Ec18 have side-chain densities, consistent with roles in the formation and recognition of the binding pocket, respectively. Among these residues, Phe318 is sandwiched by two hydrophobic residues Leu73 and Val75 in Ec18, representing another notable recognition feature.

There are two SH3 and two GK domains in the structure. The overall structure for the GK and SH3 domains are similar: the root mean square deviation (RMSD) is 1.18 Å for 205 C_α_ atoms in the GK domain and 1.23 Å for 65 C_α_ atoms in the SH3 domain. Nevertheless, we found conformational changes for two SH3 loops associated with Ec18 binding (**Fig. 2d**). One loop containing residue Leu403 moved as much as 4.5 Å; and the other loop containing Leu369 also moved so that Leu369 has a closer engagement with Leu74 from Ec18. We thus propose that residues Leu369 and Leu403 from the SH3 domain further stabilize Ec18 binding to the PDZ domain.

## Discussion

Many viruses have developed strategies to hijack human PDZ-domain containing proteins to increase their virulence and evade immune responses ^19,20^. The structure of the PSG-Ec18 complex allows us to explain the SARS-CoV-2 E-mediated PALS1 relocation and vascular damage.

PALS1 is an integral part of an apical cell polarity complex consisting of Crumbs, PALS1, and PATJ ^10^. Under physiological conditions, PALS1 interacts with the Crumbs C-terminus (Crb-CT) through the PSG module ^9^ and interacts with PATJ through its N-term L27 domain ^21^ (**Fig. 3a**). In SARS-CoV-2 infected lung epithelial cells, the replication and transcription of the virion genome produce a high load of the E protein which localizes to the ERGIC region for viral assembly and budding ^22^. The specific interactions between Ec18 and PALS1 PDZ/SH3 recruit PALS1 to the site of virus assembly and disrupt the polarity complex and vascular structure ^4^. Although the affinity between E and PALS1 is at a μM range (**Fig. S1d**) ^14^, the viral E can have a high local concentration and competes dynamically with its physiological ligand of Crumbs and pulls PALS1 out from the intercellular junction space (**Fig. 3b**). Consequently, the inter-epithelial junctions loosen and leak. The leaking junctions promote local viral spreading, flow of fluid and multiple types of immune cells (such as monocytes and neutrophils) into lung alveolar spaces, causing lung damage, and cytokine storm (**Fig. 3b**) that eventually leads to ARDS, a causative factor that contributes to the severity of symptoms and deaths in a subset of COVID-19 patients.

**Figure 3.**
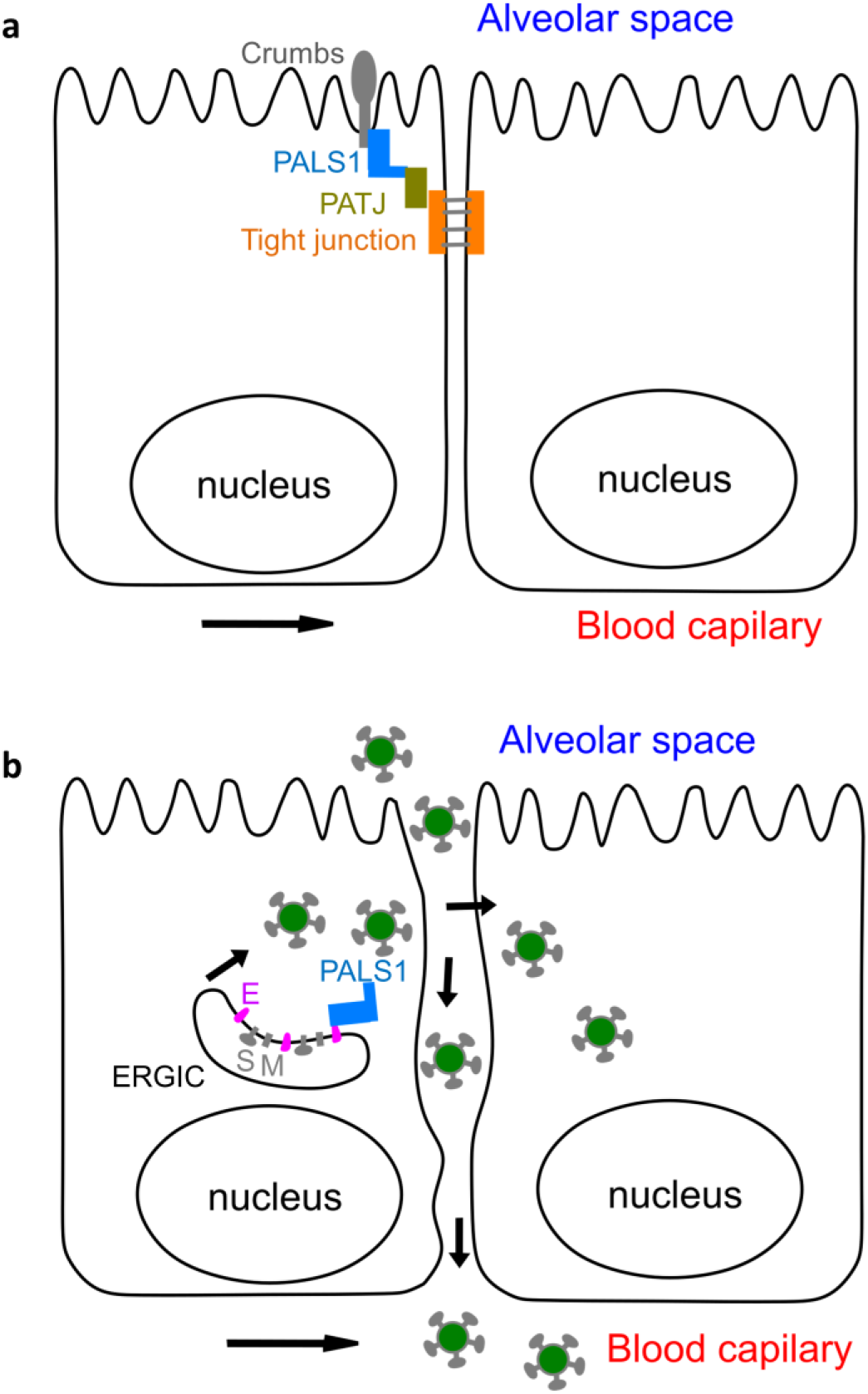
A proposed model of the E-mediated PALS1 relocalization and vascular damage. (**a**) A schematic drawing of two adjacent lung epithelial cells with the Crumbs apical complex maintaining cell polarity and tight junction formation. (**b**) SARS-CoV-2 E protein interacts with PASL1 and recruits it to the ERGIC site, causing vascular leakage and damage to cell junctions, promoting viral spreading and cytokine storm, and leading to ARDS and in some cases deaths.

Virus-host interactions have been proposed to potentiate viral fitness and virulence ^23^. Mutation in viruses, including SARS-CoV-2, that convey a selective advantage with respect to replication, assembly, release and spreading can accelerate the viral life cycle. In this work, we provide the first structure to show interactions between the SARS-CoV-2 E protein and human PALS1. The complex structure provides an atomic basis to explain E-mediated virulence through its C-terminal PBM ^13^. Interestingly, when the C-term PBM was deleted in SARS-CoV variants, the PBM was recovered from passage in the host, clearly demonstrating its involvement in viral fitness and virulence. In addition to E, SARS-CoV-2 and SARS-CoV-1 encode another PBM-containing protein ORF3a, which could both be involved in the recruitment of PDZ-containing proteins for viral fitness and virulence ^24^.

The interaction between Ec18 and PALS1 is much weaker than the interaction between PALS1 and its physiological ligand Crb-CT ^9^. Based on the alignment of PSG-Ec18 and PSG-Crb-CT structures (**Fig. S4b**), the terminal isoleucine in Crb-CT is inserted deeply in the PDZ pocket. So, a peptide inhibitor with a C-terminal isoleucine, leucine, or even phenylalanine may penetrate the pocket deeper and with a higher affinity. In addition, an arginine in Crb-CT PBM interacts with Phe318 unfavorably; changing this residue to a hydrophobic residue such as leucine or phenylalanine may enhance its hydrophobic interactions with Phe318. Interestingly, through mutations in hosts, SARS-CoV-2 variants have acquired a number of mutations in the PBM (72DLLV75) for viral fitness and virulence. Notable PBM mutations are D72Y, D72H, L73F, V75L, V75F ^25,26^. We hence propose that a hybrid peptide containing Crb-CT and viral PBM mutations could weaken the PALS1-E interactions to suppress E-mediated lung damage and virulence.

## Acknowledgments

We thank Gongrui Guo for initial discussions on the project, LBMS staff for the help with the Cryo-EM operation and data acquisition, and Computational Science Initiative (CSI) for support on computation. This work was supported in part by Brookhaven National Laboratory COVID-19 LDRD. LBMS is supported by the U.S. Department of Energy, Office of Science, Office of Biological and Environmental Research. Atomic coordinates and experimental map have been deposited in the RCSB Protein Data Bank (PDB) under the accession code XXXX.

## Author contributions

Q.L. designed the study and experiments. J.C., Y.C., C.P., L.W. and Q.L. performed the experiments. J.C., S.M., J.S., and Q.L. analyzed the data. Q.L. wrote the manuscript with help from the other coauthors.

## Methods

### Protein expression and purification

The gene encoding PASL1-PSG domains (residues 236–675) with a deletion between 411-460 was codon optimized for bacterial expression and synthesized by Genscript (www.genscript.com) and cloned into pET16-b with an N-terminal 10x his tag followed by a TEV cleavage site.

The protein was overexpressed in *Escherichia coli* BL21 (DE3) pLysS at 16°C for 18 hrs induced by addition of 0.4 mM IPTG (final) to the cell culture with an A600 of 1.0. Harvested cells were resuspended in extraction buffer containing 30 mM Tris, pH 7.5, 150 mM NaCl, 1.0 mM TCEP, 0.2 mM PMSF. Cells were lysed using an EmulsiFlex-C3 Homogenizer (Avestin, Ottawa, Canada). After centrifugation at 26,000xg for 30 mins, the supernatant was collected for affinity purification by nickel-nitrilotriacetic acid affinity chromatography (Ni-NTA, Superflow, Qiagen). The eluate was concentrated and buffer exchanged for tag removal by incubation with TEV protease overnight at 4 °C. The protein-containing solution was passed through Ni-NTA resin again to remove the cleaved tag and the protein flow-through fractions were collected, concentrated and applied to a size-exclusion column (TSKgel G3000SW column, Tosoh Bioscience) pre-equilibrated with 25 mM Tris, pH 7.5, 100 mM NaCl, 1 mM TCEP. Highly enriched protein was concentrated to about 10 mg/ml using an Amicon Ultra-15 centrifugal filter with a molecular cutoff of 30 kDa (Milipore, Inc).

### Cryo-EM sample preparation and data collection

To make the PSG-Ec18 complex, we mixed PSG and Ec18 at a molar ratio of 1:10 at a final concentration of 2 mg/ml. After incubation for 2 hrs at room temperature, we added 0.05% CHAPS (3-((3-cholamidopropyl) dimethylammonio)-1-propanesulfonate) to the sample immediately before applying 3 μl of the sample to a glow-discharged QuanFoil Au grid (0.6/1.0). Vitrification was performed using a ThermoFisher Mark IV vitrobot with a blotting condition of 3.5 sec blot time, 0 blot force, and 100% humidity at 6 °C.

Cryo-EM data were collected with the use of a ThermoFisher Titan Krios (G3i) equipped with a Gatan K3 camera and a BioQuantum energy filter. With a physical pixel size of 0.684 Å, a total dose of 64 e^−^/Å^2^ were fractioned to 52 frames under the super-resolution mode using the ThermoFisher data acquisition program EPU. A total of 12,861 movies were collected with an energy filter width of 20 eV throughout the data acquisition. Data collection statistics is listed in **Table S1**.

### Cryo-EM data analysis

Beam-induced motion correction was performed using MotionCorr2 ^27^ through a wrapper in Relion3 ^17^ with a bin-factor of 2. Corrected and averaged micrographs were further corrected by CTF estimation using Gctf ^28^. Micrographs with an estimated resolution lower than 4.5 Å were discarded from further processing. Particle picking was performed using Localpicker ^16^ which uses per-micrograph defocus values (estimated by Gctf) to set up picking parameters. We picked up a total of 6,375,890 particles, extracted and binned them to 64 pixels with a pixel size of 2.736 Å.

We used CryoSPARC ^18^ and Relion3 for 2D and 3D class averages and 3D refinements. Specifically, we used 2D class averaging for initial particle cleanup which resulted in 2,193,282 particles. Among these particles, we produced an initial 3D model using CryoSPARC and used the model to perform 3D classifications in Relion3 for five classes with a pixel size of 2.736 Å (**Fig. S2**). Particles from the 3D class with the best structural feature were selected. A total of 715,010 particles were selected, re-centered, and re-extracted at 256 pixels with a pixel size of 0.684 Å.

Extracted particles were further auto-refined to convergence with Relion3 followed by non-alignment 3D classification into three classes (**Fig. S2**). Particles from the best class (7.2%) were selected for CTF refinement and Bayesian polishing in Relion3 and non-uniform refinement in CryoSPAC to reach a refined reconstruction at 3.65 Å resolution based on gold-standard Fourier Shell Correlation of 0.143 (**Fig. 1d**). Local resolutions were estimated using BlocRes ^29^. Reconstruction statistics is listed in **Table S1**.

### Model building and refinement

To assist our model building and refinement, we sharpened the masked and filtered map using PHENIX ^30^ with a B factor of −100 Å^2^. We used the PDB code 4WSI as a starting model and built the model for PSG and Ec18 in COOT ^31^ and refined the model iteratively using PHENIX. The refined model was validated using Molprobity ^32^ and the refinement statistics is listed in **Table S1**.

### Microscope thermophoresis (MST) measurement

The binding affinity between Ec18 and PALS1-PSG was measured using a Monilith NT.115 instrument (Nanotempertech). Purified protein was buffer exchanged and covalently labeled using dye NT647 following manufacture’s protocol. The labeled protein was diluted 10x prior to measurement in assay buffer containing 50 mM Tris, pH 7.5, 100 mM NaCl, 1 mM DTT, 1 mM EDTA, and 0.05% Tween 20. Ec18 was dissolved in the assay buffer to a final concentration of 5.0 mM. Ten microliters of Ec18 was diluted 1:1 serially in the assay buffer and mixed with an equal volume of labeled PSG. The PSG-Ec18 samples were incubated for 10min in dark at room temperature before MST measurements. For all MST measurements, we used a MST power medium, laser power 40%, and MST time 30 sec. NanoTemper program MO.Affinity Analysis (Nanotempertech) was used for data analysis and curve fitting with a K*d* model.

---

**Extended Data Fig. 1.**
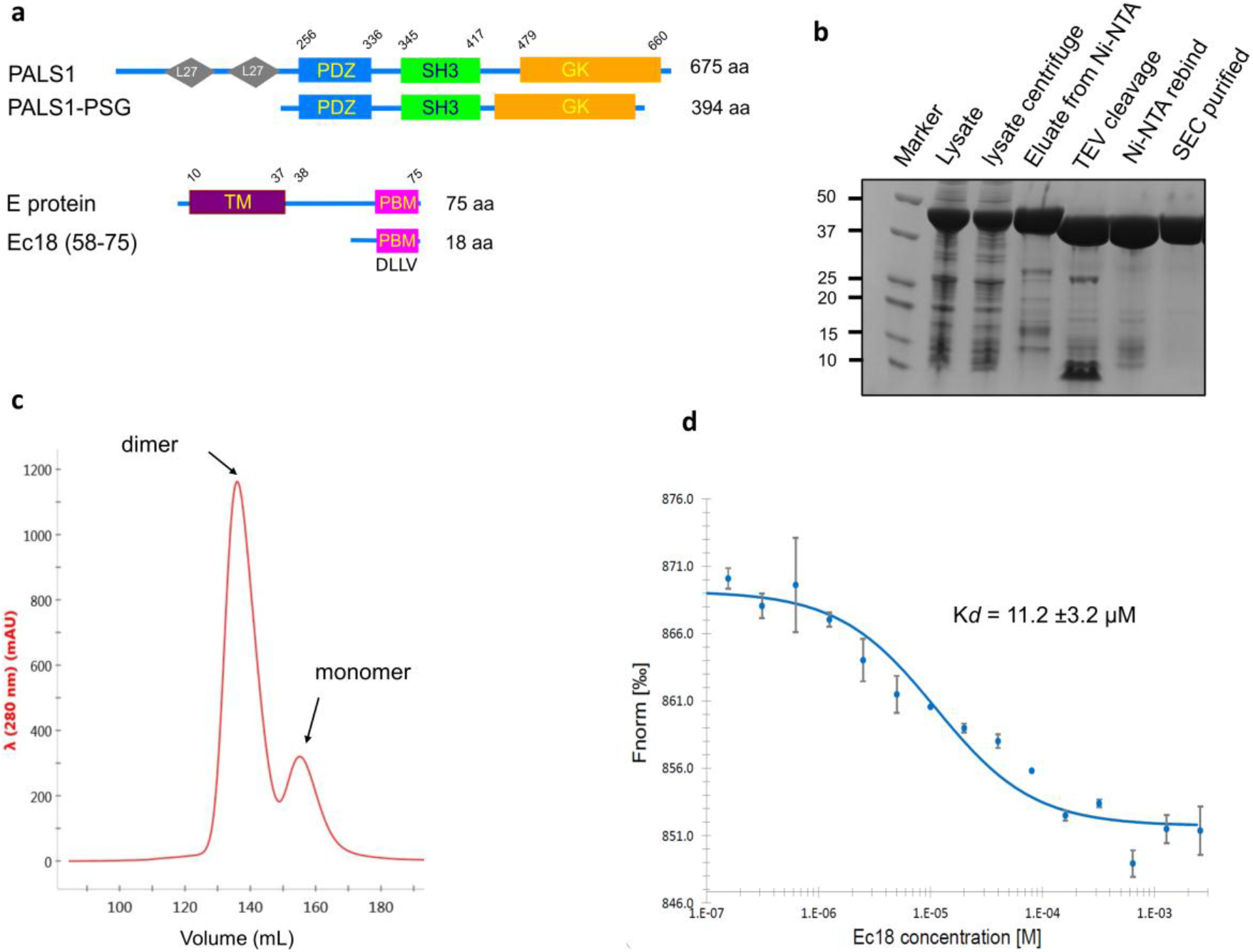
Protein production. (**a**) Schematics of the domains in PALS1 and E. PALS1-PSG and Ec18 were used in this work. (**b**) SDS-PAGE analysis for the purification of PSG. (**c**) SEC analysis for the purification of the PSG dimer. (**d**) Interactions between PALS1-PSG and Ec18 measured by microscale thermophoresis (MST). PALS1-PSG was labeled by a fluorescence dye NT-647 and was titrated using Ec18 at different concentrations. The fitted K*d* is 11.2 ±3.2 μM. The error bar is the standard deviation from experiments of two different samples.

**Extended Data Fig. 2.**
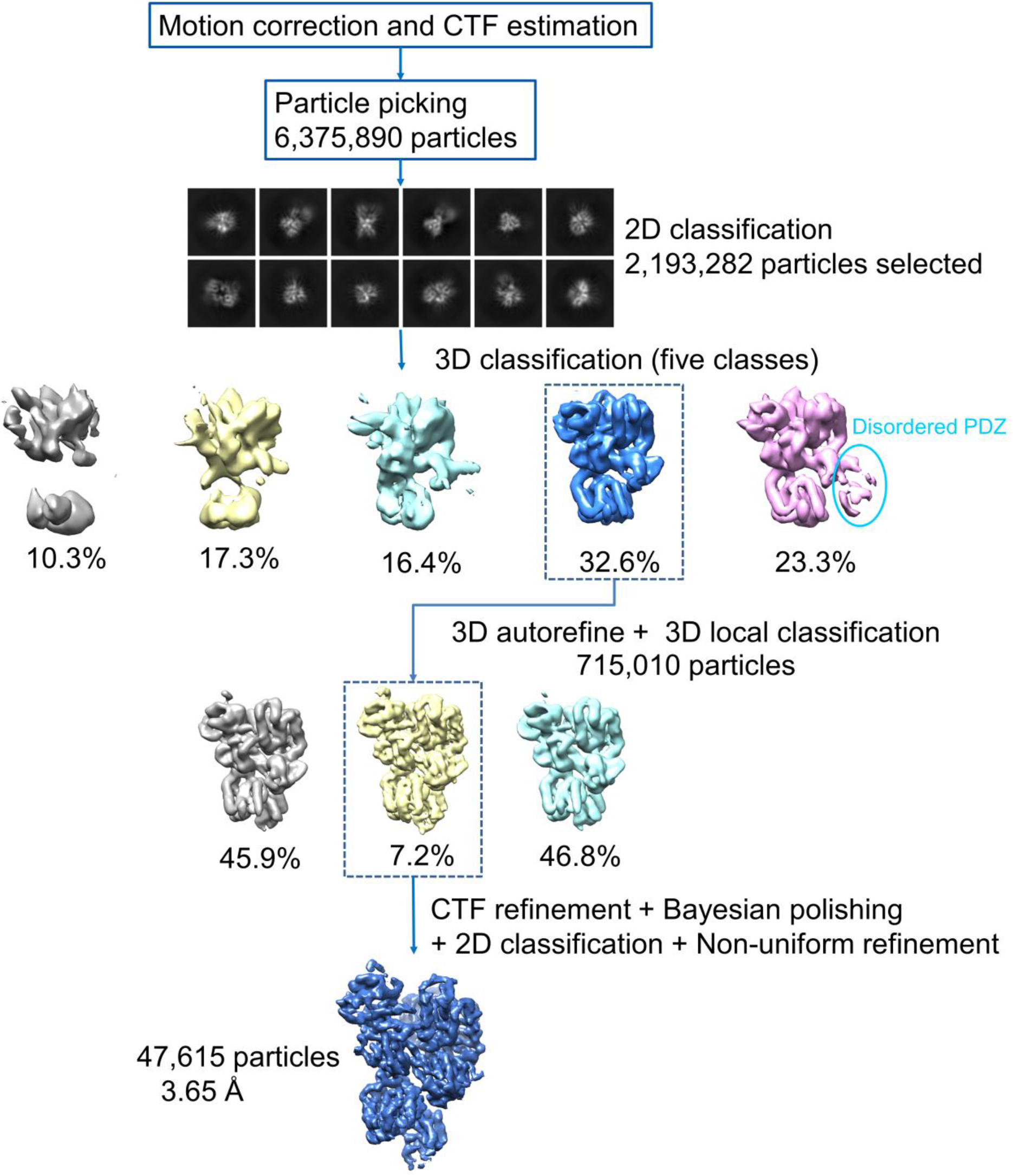
A data analysis workflow.

**Extended Data Fig. 3.**
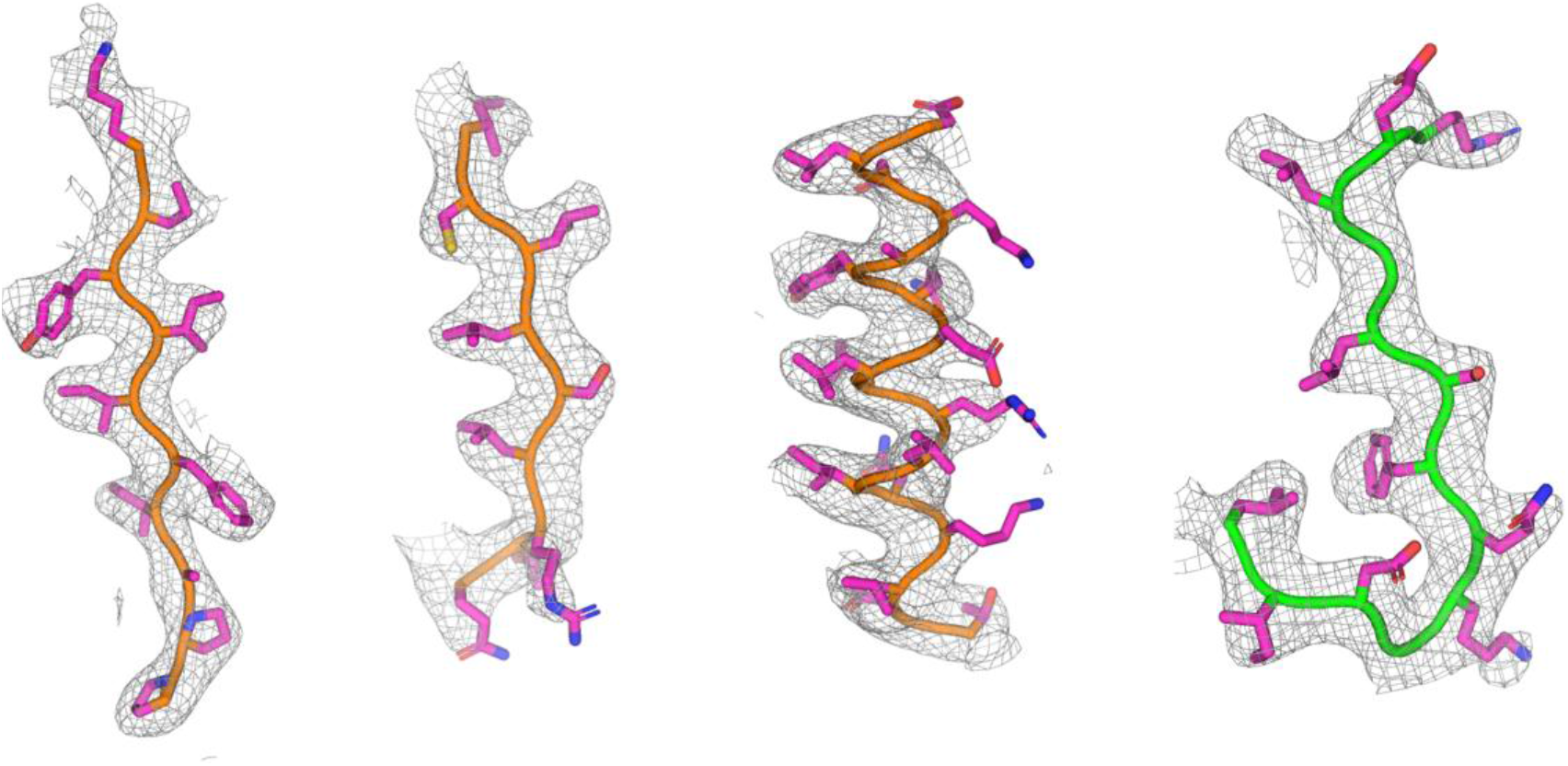
Quality of densities showing secondary structures and side chains.

**Extended Data Fig. 4.**
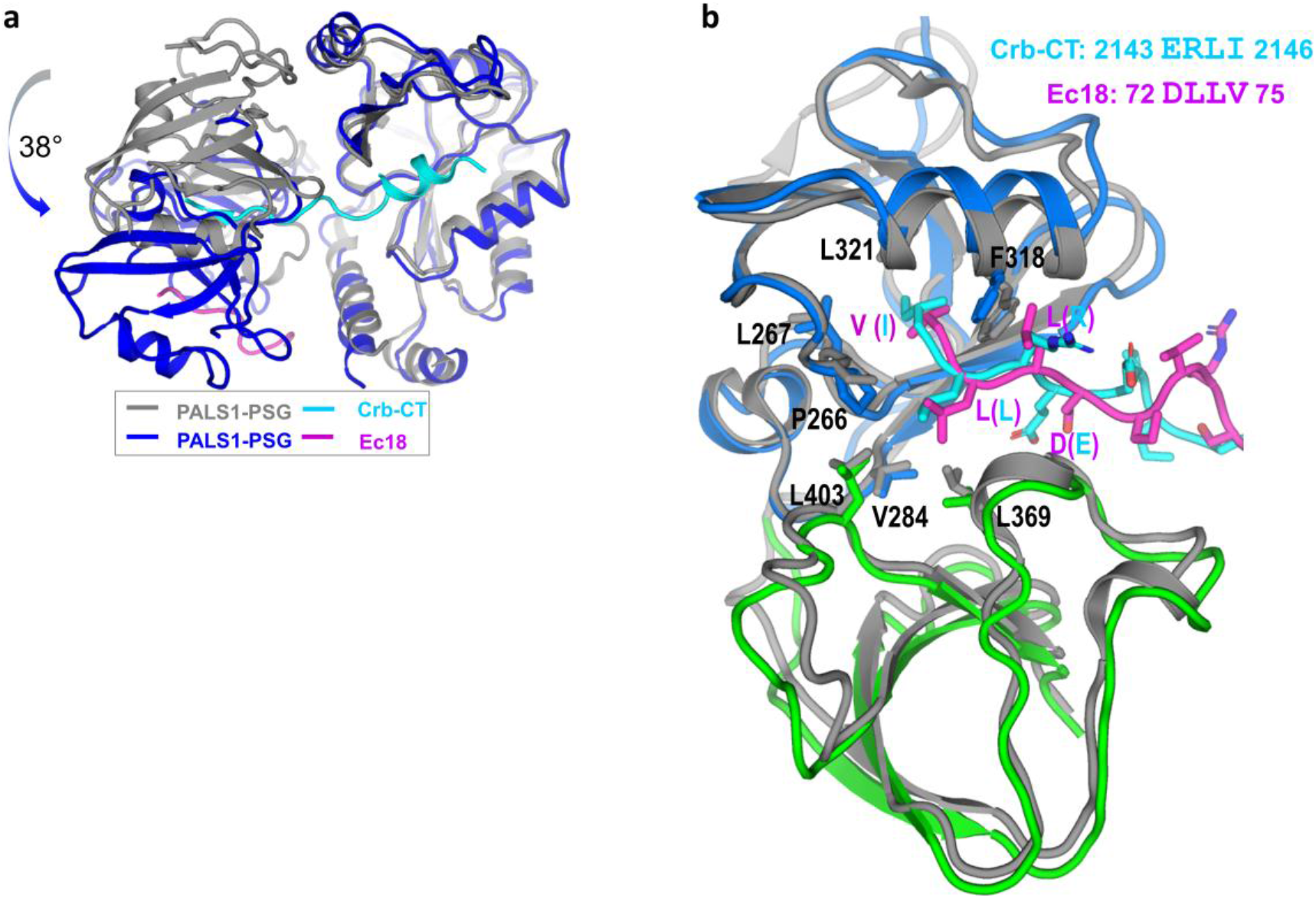
Structural comparison of PSG-Ec18 with PSG-Crb-CT (PDB code 4WSI). (**a**) Alignment of the two complex structures based on the GK domain showing a relative rotation of about 38° for the SH3-PDZ domain. (**b**) Structural superposition for the SH3 and PDZ domains showing the binding sites for Ec18 and Crb-CT.

**Extended Data Table 1.**
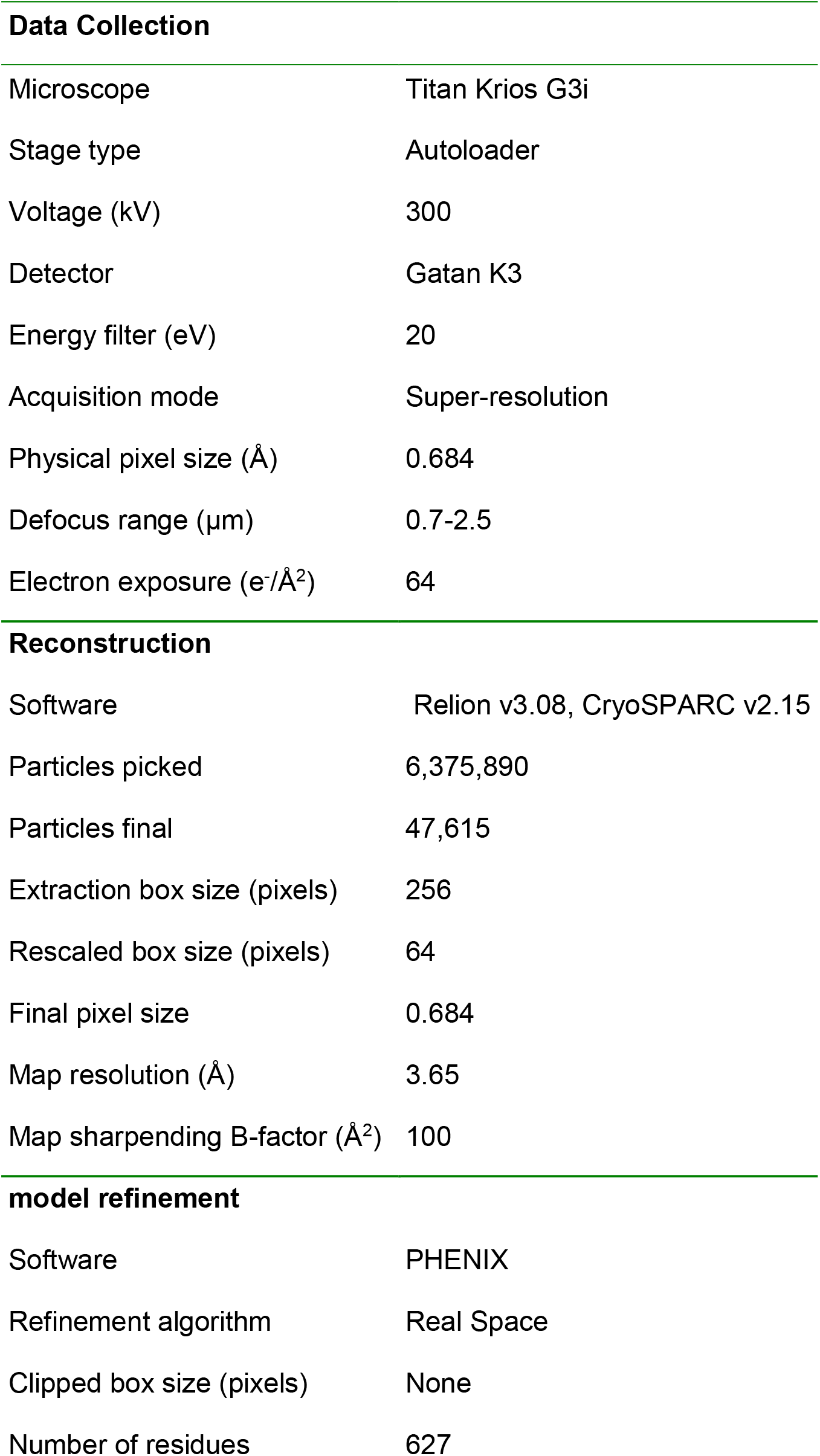
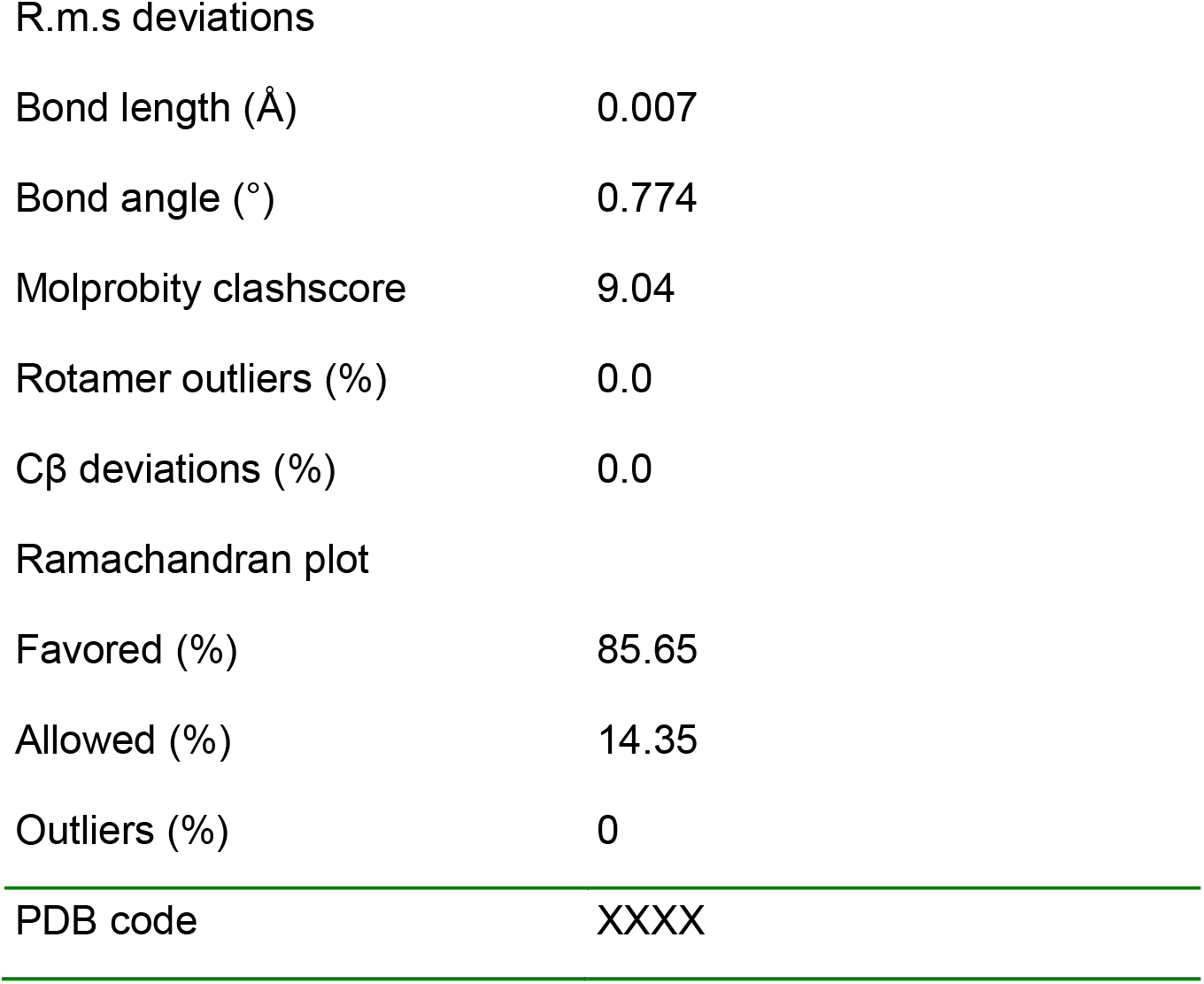
Cryo-EM data collection, 3D reconstruction, and refinement statistics.

